# Evidence for a RIPK1-independent survival mechanism for CASPASE-8 in αβ T cells

**DOI:** 10.1101/2025.03.27.645705

**Authors:** Farjana Islam, Scott Layzell, Ines Boal-Carvalho, Benedict Seddon

**Affiliations:** Institute of Immunity and Transplantation, Division of Infection and Immunity, University College London, The Pears Building, Hampstead, London, UK; Department of Biochemistry and Molecular Biology, Shahjalal University of Science and Technology, Sylhet 3114, Bangladesh; The Breast Cancer Now Toby Robins Research Centre, The Institute of Cancer Research, London, UK

## Abstract

CASPASE8 promotes both cell death and survival by acting as a trigger of apoptosis but also a repressor of necroptosis. In T cells, CASPASE8 is required for FAS induced apoptosis, but protects activated T cells from necroptosis. The broader role and mechanisms of CASPASE8 in the wider T cell compartment is less certain. Here, we analysed mice in which *Casp8* was conditionally deleted in the lymphoid compartment by huCD2iCre. In the thymus, we found evidence of a modest impairment of early thymic progenitors and a striking absence of NKT cell development. Amongst mature peripheral T cells, there was a substantial and specific reduction in the CD8 T cell compartment, that included naive, central memory and virtual memory subsets. While life spans and turnover of CD8 T cells appeared largely normal, we did identify an acute requirement for continued CASPASE8 expression for survival of a fraction of CD8 T cells, since induced *Casp8* deletion by tamoxifen induced CD8^CreERT^ resulted in an acute loss of CD8 T cells. CASPASE8 deficient T cells were resistant to FAS or TNF induced cell death in vitro. Generating *Casp8* deficient mice that express a kinase dead RIPK1 confirmed that necroptosis contributed to death of thymic progenitors and some peripheral CD8 T cell subsets in the absence of CASPASE8. However, kinase dead RIPK1 failed to restore NKT cell development and only partially rescued CD8 VM T cells, while analysing mixed bone marrow chimeras suggested that CASPASE8 deficient CD4 and CD8 T cells were less competitively fit than WT T cells, even in the absence of RIPK1 kinase activity. These latter observations suggest the existence of a novel mechanism by which CASPASE8 promotes T cell survival that is independent of its established role in repressing necroptosis.

## Introduction

CASPASE-8 is a critical regulator of extrinsic cell death pathways. Receptor triggered recruitment of CASPASE8, mediated by the adapter FADD, results in oligomerisation of pro-CASPASE8, that leads to proximity-induced autocatalytic cleavage (Tummers and Green, 2017). TNFR superfamily member containing death domains such as Fas (CD95/Apo-1), DR3 (Apo-3), DR4 (Apo-2 or TRAILR1), DR5 (TRAILR2) trimerise upon ligand binding and can directly recruit FADD (Ashkenazi and Dixit, 1998). In contrast, TNFR1 recruits FADD via the adapter TRADD, but only triggers cell death under conditions in which the initial signalling complex-I becomes destabilised (Annibaldi and Meier, 2018). Other receptors capable of recruiting FADD include Toll-like receptors (Zhande et al., 2007), NLRP3 and other inflammasomes (Mouasni et al., 2019), Dectin-1 (Ketelut-Carneiro et al., 2018) and type I interferon receptors (Thapa et al., 2013). Once cleaved, active CASPASE8 is released into the cytoplasm which cleaves and activates executioner caspases (CASPASE3, CASPASE7). In some cells such as hepatocytes and fibroblasts, CASPASE-8 cleaves Bid to tBid, which promotes the release of Second Mitochondria-Derived Activator of Caspases (SMAC) into cytosol that inhibits IAPs and triggers apoptosis (Li et al., 1998;Yin et al., 1999). As well as inducing cell death, CASPASE-8 actively inhibits necroptosis. In the absence of CASPASE8, RIPK1 can recruit RIPK3 which in turn activates the pore forming capacity of MLKL resulting in necroptosis (Tenev et al., 2011). CASPASE8 is thought to target cleavage of RIPK1 thereby preventing necrosome formation (Newton et al., 2019;Newton et al., 2024). As such, CASPASE8 is a critical regulator of cell death and cell survival. Mice lacking CASPASE8 expression die in utero at E10.5 as a result of uncontrolled necroptotic cell death, resulting in vascular and haematopoetic defects, that is rescued by additional deletion of RIPK3 (Kaiser et al., 2011).

The role of CASPASE8 in T cells has been studied using a variety of conditional gene deletion models, all of which suggest a critical role of CASPASE8 for normal homeostasis of the T cell compartment, but the details of which vary depending on *Cre* driver used. Studies in which deletion is mediated by Lck-Cre report normal thymic development, but profound peripheral T cell lymphopenia, even though T cells are resistant to FAS induced cell death (Salmena et al., 2003). The same group later report that mice develop a lymphoproliferative disorder that had features resemblent of FAS deficient *Fas^lpr^* mice, consistent with a loss of CASPASE8 dependent FAS triggered cell death (Salmena and Hakem, 2005). Lck-Cre has subsequently been shown to induce significant Cre toxicity (Shi and Petrie, 2012) which may be a compounding factor in the phenotype exhibited by this strain, and could account for the reduction in peripheral T cell numbers. Studies employing CD4^Cre^, that does not induce such Cre toxicity (Shi and Petrie, 2012), report normal numbers of thymocytes and peripheral T cells in lymph nodes and spleen (Ch’en et al., 2008) though later studies suggest a subtle reduction in percentages of T cells in lymph nodes and a reduction in the frequency of CD8^+^ T cells, though not significant (Ch’en et al., 2011). Activation of CASPASE8 deficient T cells, either in vitro by CD3 crosslinking or in vivo following infection with LCMV, resulted in defective responses (Ch’en et al., 2008) that could be rescued either by blocking RIPK1 kinase activity or RIPK3 deletion (Ch’en et al., 2011), revealing the role of CASPASE8 in blocking necroptotic cell death in activated T cells. Mice with CD4^Cre^ mediated *Casp8* deletion and with additional loss of RIPK3 resemble *Fas^lpr^* mice, since they developed expansions of CD4^−^CD8^−^ double negative T cells (Ch’en et al., 2011). While CASPASE8 blocks necroptotic cell death of acutely activated T cells, it is unclear whether resting naive or memory phenotype cells are susceptible to necroptosis in the steady state. CD44^hi^ memory phenotype CD4 and CD8 T cells are reportedly normal in the absence of CASPASE8 (Ch’en et al., 2008) and it was not reported whether RIPK3 deficiency rescues the modest reduction in T cell frequencies described in later studies (Ch’en et al., 2011). In contrast, induction of necroptotic cell death amongst Foxp3^+^ regulatory CD4^+^ T cells has also been demonstrated using Foxp3^Cre^ mediated deletion of *Casp8* (Teh et al., 2022), and results in immune perturbations expected from a loss of regulatory cell activity, that is rescued by additional MLKL ablation.

In the present study, we wished to resolve the apparent discrepancies in the literature around the role of *Casp8* for normal homeostasis of peripheral αβ T cell compartments in the steady state and provide a deeper analysis of mature αβ T cell subsets. Using huCD2^iCre^ driver line, that deletes *Casp8* from early double negative thymic progenitors, and does not induce Cre toxicity (Shi and Petrie, 2012), we find evidence that CASPASE8 is required for normal thymopoiesis of αβ T cells and in various peripheral mature T cell populations. However, using kinase dead RIPK1 to block necroptotic cell death, we find that some but not all of the defects resulting from *Casp8* deletion can be rescued suggesting a prosurvival function of CASPASE8 that is distinct from its recognised function in repressing necroptosis.

## Results

### CASPASE8 expression is required for optimal expansion of DP thymic compartment but not selection or development of SPs

To assess the impact of *Casp8* deletion on thymic development, we first analysed the thymic phenotype in *Casp8^fx/fx^* huCD2^iCre^ mice (Casp8ΔT^CD2^) as compared with Cre -ve littermates. The distribution of subsets defined by CD4 and CD8 expression was largely normal. A modest but significant reduction of total thymus size was evident in the absence of CASPASE8, which was reflected in a reduction in both CD4^−^CD8^−^ double negative (DN) and CD4^+^CD8^+^ double positive (DP) compartments (Fig. 1A). Analysing the size of precursor DN1-4 subsets, as defined by CD44 and CD25 expression, revealed normal DN3 numbers but a reduced DN4 compartment (Fig. 1B). In contrast, other aspects of conventional thymic development appeared normal. CD4 and CD8 single positive (SP) compartments were largely normal in size and maturation within these compartments also appeared normal (Fig. 1C and 1D), suggesting the loss in DP cellularity was not sufficient to adversely impact the size of mature post-selection compartments and therefore thymic output. However, analysing the development of NKT cells, identified by their binding to CD1d tetramers complexed with glycolipid α-galactosylceramide (CD1d-α-GalCer), revealed a profound defect in NKT cells within the thymus. This appeared to be associated with a failure of mature CD44^hi^ NKT cells to accumulate, since CD1d-α-GalCer binding precursors amongst DP thymocytes appeared normal as were immature HSA^hi^ precursors amongst DN and CD4 SP populations (Benlagha et al., 2005) (Fig. 1E).

**Figure 1.**
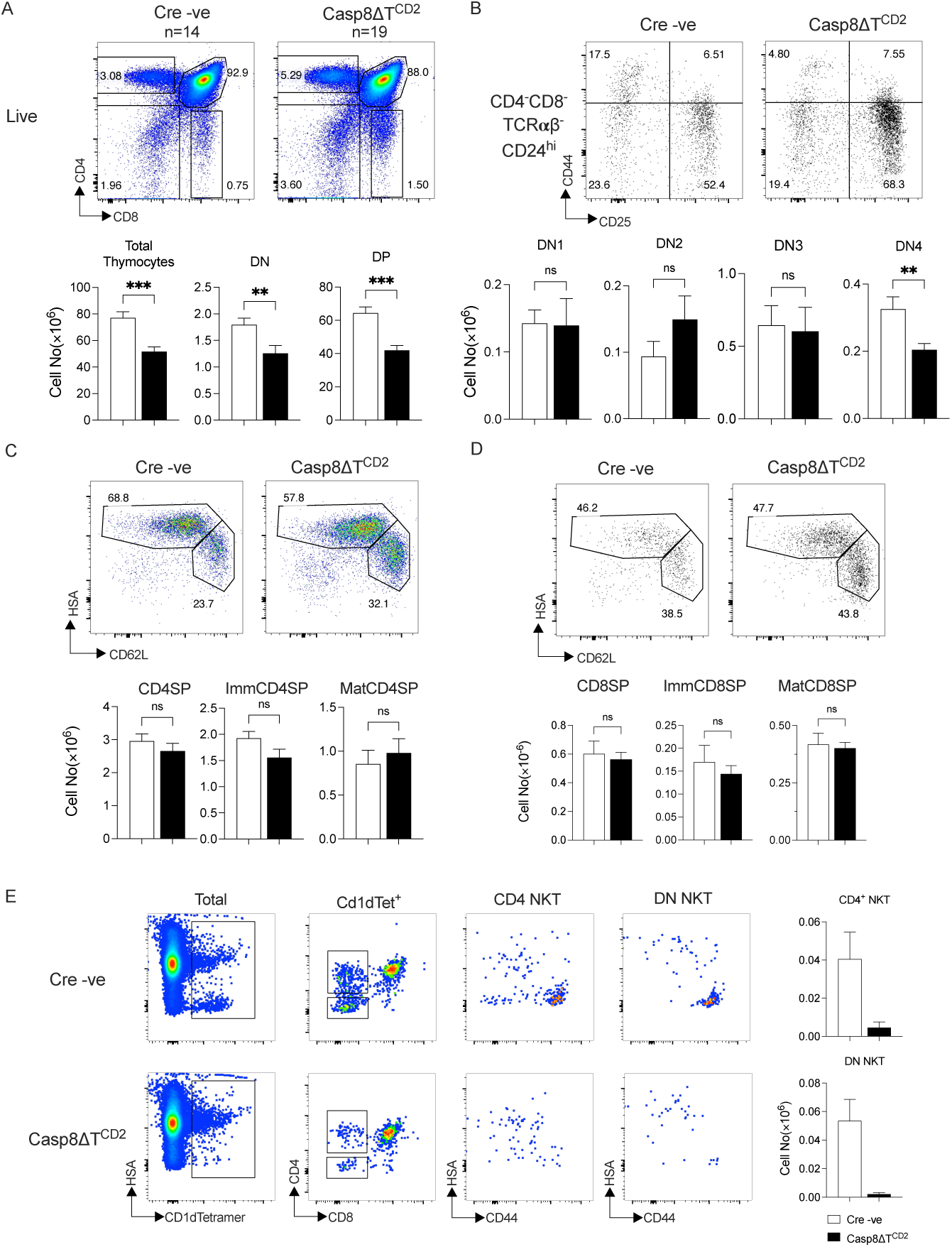
CASPASE8 deficiency impacts αβ T cell development. Thymi from Casp8ΔT^CD2^ mice (n=19) and Cre -ve litter mates (n=14) were enumerated and analysed by flow. (A) Density plots are representative expression of CD4 vs CD8 by live gated thymocytes. Bar charts show the numbers of total thymocytes, CD4^−^CD8^−^ DN thymocytes and CD4^+^CD8^+^ DP thymocytes. (B) Dot plots are of CD44 vs CD25 expression by CD4^−^CD8^−^TCRb^−^ HSA^hi^ thymocytes from representative mice. Bar charts summarise numbers of different DN subsets. (C-D) Density and dot plots of HSA vs CD62L are from CD4^+^CD8^−^ TCRβ^+^ (C) and CD4^−^CD8^+^ TCRβ^+^ (D) live thymocytes. Bar charts summarise the total numbers of SP, immature HSA^hi^CD62L^lo^ and mature HSA^lo^CD62L^hi^ SP subsets. (E) Density plots are of HSA vs CD1d-α-GalCer (CD1dTet) binding to total live thymocytes. CD4 vs CD8 expression is for CD1dTet^+^ cells. HSA vs CD44 expression is shown for CD4^+^CD8^−^ CD1dTet^+^ (CD4 NKT) cells and CD4^−^CD8^−^ CD1dTet^+^ (DN NKT) cells. Bar charts summarise total numbers of CD4 NKT and DN NKT recovered from mice. Data are pooled from multiple batches of mice analysed.

### Perturbations of CD8 T cell subsets in the absence of CASPASE8 expression

We next analysed the size and detailed composition of peripheral compartments of CD4 and CD8 T cells in lymph nodes and spleen, to determine the impact of *Casp8* deletion. Analysing the CD4 T cell compartment revealed relatively normal size and composition, except that CD4 effector memory compartment appeared slightly enlarged, relative to other subsets (Fig. 2A), though this was not reflected by a detectable change in absolute cell numbers (Fig. 2B). In contrast, the CD8 compartment exhibited multiple defects. Numbers of naive CD44^lo^, CD62L^hi^ CD49d^hi^ CD44^hi^ central memory (CM) and CD62L^hi^ CD49d^lo^ CD44^hi^ virtual memory (VM) subsets were all reduced. In contrast, absolutely numbers of CD62L^lo^ CD44^hi^ effector memory (EM) CD8 cells were similar to controls, even though other subsets were reduced, and this was reflected by an increased representation of these cells amongst total CD8^+^ T cells (Fig. 2B). A reduction in the sized of the CD8 compartment could result from several causes, including reduced self-renewing cell divisions. To assess division as a potential cause of the defects observed in the CD8 compartment we assessed expression of the nuclear antigen Ki67 that is expressed by cells during mitosis. We assessed division both amongst SP subsets in the thymus, that undergo limited division before egress (Rane et al., 2022), and amongst naive, CM, EM and VM subsets. In general, no differences in Ki67 expression were observed, with the exception of CD8 VM cells, and a small increase amongst CD8 naive cells, that typically express very low levels of Ki67 (Fig. 2C, D). Therefore we found no evidence that the reduction in CD8 subsets was due to a failure of self-renewing cell division processes.

**Figure 2.**
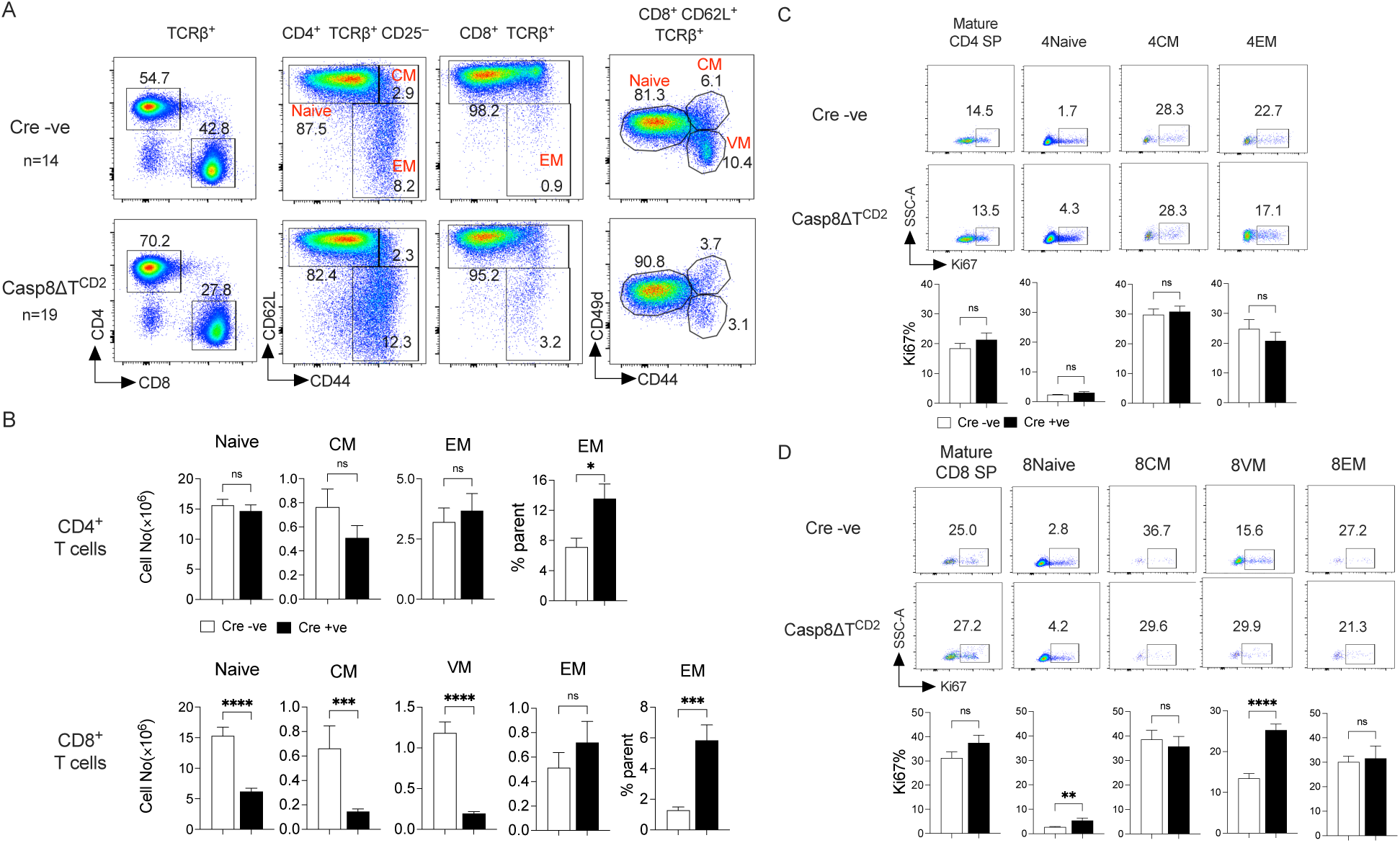
Reduced peripheral CD8 compartment size following *Casp8* deletion. Lymph nodes and spleen from the cohorts of mice described in figure 1 were enumerated and analysed by flow. (A) Density plots are of CD4 vs CD8 by TCRβ^+^ lymph node cells, CD62L vs CD44 by either CD4^+^ TCRβ^+^ CD25^−^ or CD8^+^ TCRβ^+^ subsets. Gates used to identify CD4^+^ naive, CM and EM subsets are labelled in red. Naive, CM and VM CD8 subsets were identified amongst CD8^+^ CD62L^hi^ TCRβ^+^ cells using CD49d and CD44 as indicated. (B) Bar charts summarised the total numbers of the indicated CD4 and CD8 subsets from total lymph nodes and spleen combined. (C-D) Density plots are of SSC vs Ki67 by the indicated subsets of CD4 and CD8 T cells in thymus and lymph nodes. Bar charts summarise the frequency of Ki67^+^ cells amongst the indicated subsets.

### Normal survival signalling and dynamics in the absence of CASPASE8

We next assessed survival signalling amongst naive T cells. TCR induced survival signals are required for optimal naive T cell survival, and can be assessed by measuring the surrogate marker CD5 (Seddon and Zamoyska, 2002). CD5 expression appeared at normal levels in both CD4 and CD8 naive T cells from Casp8ΔT^CD2^ mice implying similar extents of TCR signalling (Fig. 3A). IL-7 is also an essential survival cue for naive T cells, and IL-7 signalling is regulated in T cells at the level of IL-7R expression (Barata et al., 2019). We did observe a modest reproducible and significant reduction in the levels of IL-7R protein expressed by both CD4 and CD8 naive T cells (Fig. 3B). This reduction in IL-7R could potentially result in reduced life span of naive T cells that could account for altered compartment sizes, although we noted that CD4 naive T cell numbers were normal, in spite of reduced IL-7R expression. Nevertheless, we sought to assess survival of naive T cells from Casp8ΔT^CD2^ mice to determine whether a population wide survival defect could account for the reduction in the CD8 T cell compartment. To do this, we isolated total T cells from Cre + and Cre -ve littermates and transferred them to congenic CD45.1^+^CD45.2^+^ hosts. As further control, we also co-transferred WT T cells from CD45.1 donors with both Cre -ve and Cre +ve donor cells (Fig. 3C). Hosts were periodically bled and representation of donors T cell subsets assessed together with IL-7R expression level. As expected, control CD45.1 and Cre -ve donor CD4 and CD8 naive T cells steadily declined in number over time for both subsets, in line with previous studies and expected life spans (Seddon and Yates, 2018). Analysing survival of T cells from Casp8ΔT^CD2^ donors revealed near identical profiles of loss as exhibited by control T cells (Fig. 3D). Representation of CD8^+^ naive T cells from Casp8ΔT^CD2^ donors started from a lower point since they were reduced in numbers in the original donors. However, their decay thereafter tracked that of control populations. Analysing IL-7R expression showed that the reduced expression level amongst Casp8ΔT^CD2^ donor T cells was preserved following transfer. However, in all cases, the measured level of IL-7R on surviving T cells gradually increased over time. Therefore, we found no evidence that the lifespan of naive CD8 T cells in Casp8ΔT^CD2^ mice was reduced compared with controls, and the small difference in IL-7R expression we observed did not appear to alter life span.

**Figure 3.**
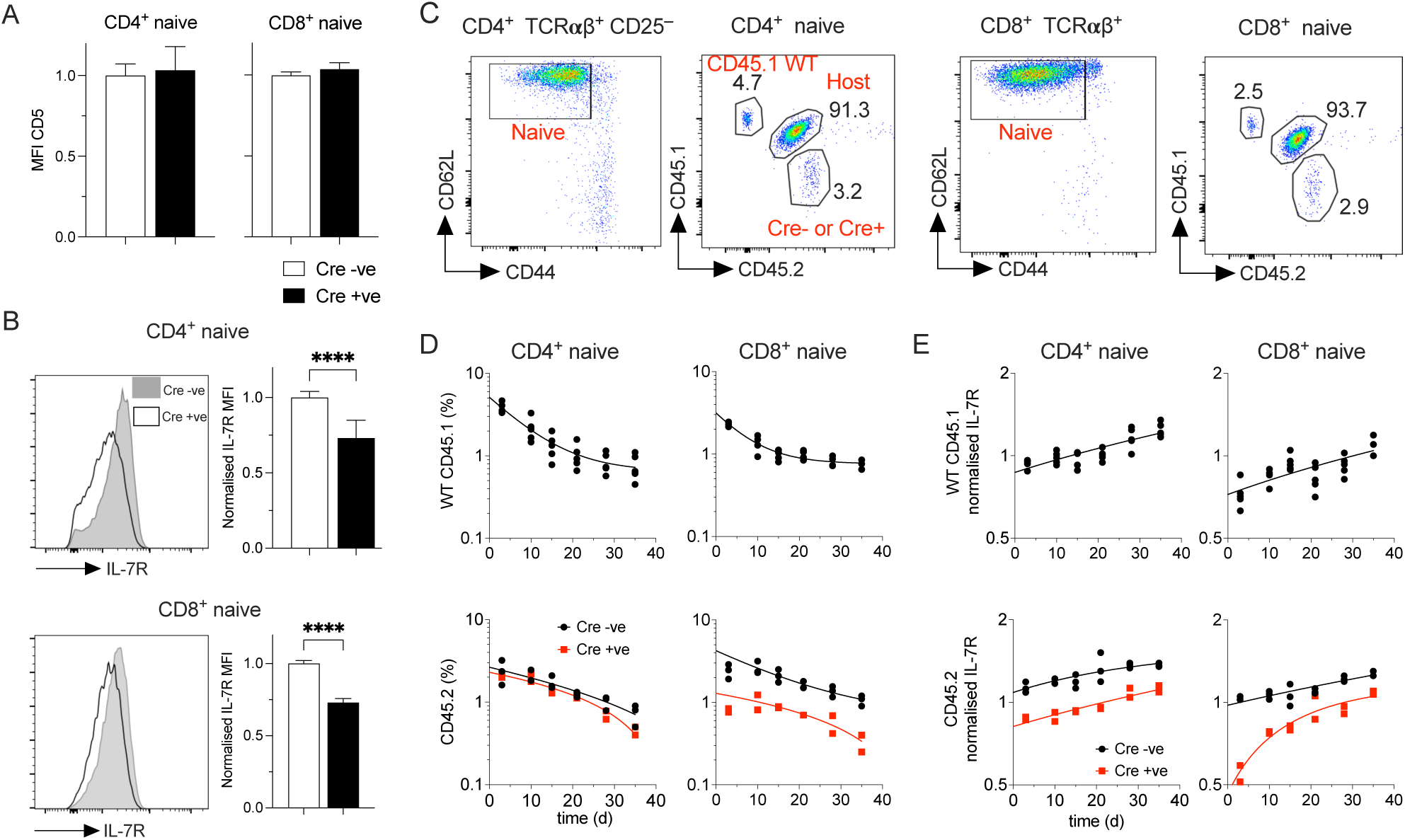
Normal survival signaling and dynamics of T cells from Casp8ΔT^CD2^ donors. (A) Bar charts are MFI of CD5 expression normalised to the average level in Cre -ve T cells of the indicated naive subsets. (B) Histograms are of IL-7R expression by naive CD4 and CD8 T cells from the indicated donors, and bar charts are summary of expression normalised to average MFI of Cre -ve controls. (C) Gating strategy to track the indicated donor populations in CD45.1^+^CD45.2^+^ hosts amongst naive CD4 and naive CD8 populations in blood. (D) Line graphs summarise representation of donor populations amongst total naive CD4 and naive CD8 populations in blood, and (E) their expression of IL-7R, normalised to average MFI of host subsets. Data are representative of two independent experiments.

### Continued CASPASE8 expression is essential for naive T cell survival

Our adoptive transfer experiments suggested that life span of T cells recovered from Casp8ΔT^CD2^ mice was comparable to that of T cells from control mice. However, the cells we assessed were those that remained in adult Casp8ΔT^CD2^ mice and may represent a subpopulation that had been subjected to counter selection over the life of the mouse, thus enriching for longer-lived cells at the expense of shorter-lived ones. This would reconcile the observations of normal lifespans but reduced compartment size, specifically for CD8 T cells. If true, this would predict that a substantial fraction of the normal peripheral CD8 T cell compartment should be reliant on continued CAPSASE8 expression for their normal survival. To test this, we bred *Casp8^fx^* mice with CD8^CreERT^ transgenic mice that express a tamoxifen inducible CreERT2 construct under control of the CD8 E8i promotor, that specifically targets expression to peripheral CD8^+^ T cells, but not thymocytes (CD8^CreERT^) (Andrews et al., 2021). It has not previously been reported whether this Cre driver exhibits any Cre toxicity. Therefore we first tested the impact of our tamoxifen treatment regime of five daily injections into hemizygous CD8^CreERT^ mice. We quantified CD8 T cell numbers at d7, to assess acute Cre toxicity, and also at d21, which is a time point when we would assay the impact of induced *Casp8* deletion. Numbers of CD8 naive and CD8 CM were normal at both d7 and d21 after first tamoxifen injection. CD8 EM were reduced by around half while CD8 VM were reduced by a third, suggesting that there was some limited *Cre* toxicity in these subsets (Fig. S1). Since naive and CM subsets were not susceptible to *Cre* toxicity in this strain, we went on to assess the impact upon these subsets of *Casp8* deletion in *Casp8^fx^* CD8^CreERT^ mice given five injections of TAM on consecutive days. At d21 after the first injection, the CD8 T cell compartments of treated mice were analysed. This revealed that both CD8 naive and CD8 CM numbers were reduced by around half (Fig. 4A-B). EM and VM populations were also reduced, but in line with reductions observed in TAM treated CD8^CreERT^ single transgenic mice (Fig. 4B). In contrast to CD8 T cells from Casp8ΔT^CD2^ mice, IL-7R levels were not altered on CD8 T cells following *Casp8* deletion (Fig 4C).

**Figure 4.**
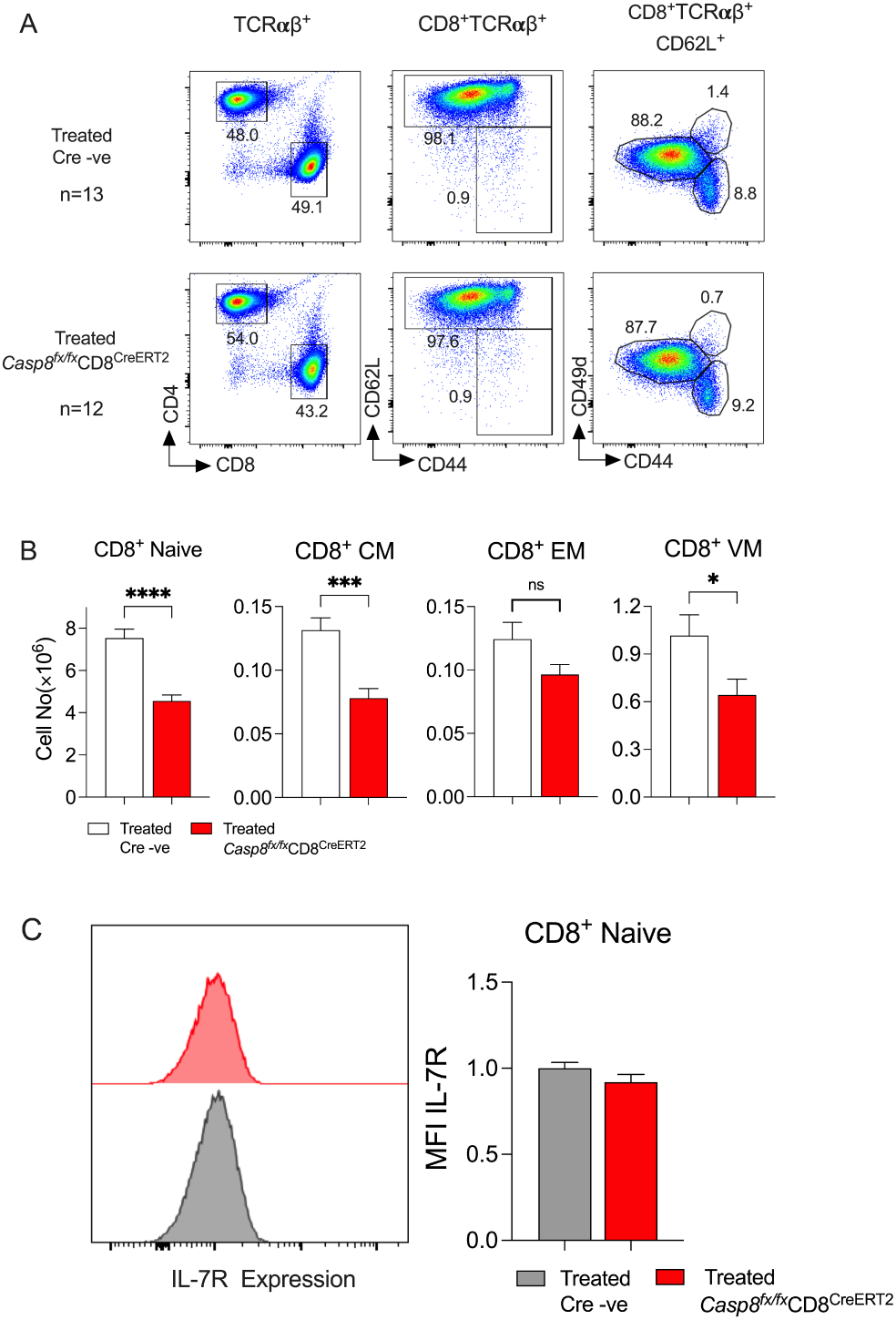
CD8 T cells require constitutive CASPASE8 expression for their survival. Casp8*^fx^* CD8^CreERT^ mice were treated with TAM for five consecutive days. At day 21, mice were culled and CD8 T cell compartment in lymph node and spleen enumerated. (A) Density plots show representative phenotypes in lymph nodes cells from TAM treated *Casp8^fx^* CD8^CreERT^ mice (n=13) and Cre-ve littermate controls (n=12) and gates used to identify naive, EM, CM and VM subsets of CD8 T cells. (B) Bar charts show total numbers of the indicated CD8 subset recovered from lymph node and spleen of the indicated strains. (C) Histograms are of IL-7R expression by naive CD8 T cells from the indicated mice. Data are pool of 4 independent experiments.

### CASPASE8 deficient T cells are resistant to both TNF and FAS induced cell death in vitro

The loss of naive T cells following deletion of *Casp8* suggested that CASPASE8 deficient T cells were susceptible to cell death by some mechanism. The proteolytic activity of CASPASE8 is known to repress necroptotic cell death by targeting RIPK1 (Newton et al., 2019) and thereby prevent formation of the necroptosome. In T cells, this activity of CASPASE8 has been described in activated T cells but not resting subsets of naive or memory T cells, which are not thought to express MLKL. Nevertheless, we wished to test whether necroptotic cell death in the absence of CASPASE8 expression could account for the loss of peripheral naive CD8 T cells. We first tested whether CASPASE8 deficient T cells were sensitive to death induced by TNFRSF members. We challenged CASPASE8 deficient T cells with two death inducing stimuli - FAS stimulation with recombinant FLAG-FASL protein and additional anti-FLAG crosslinker, and TNF in the presence of panIKK inhibitor. In the latter case, IKK inhibition sensitises T cells to RIPK1 dependent cell death when stimulated with TNF (Carty et al., 2023). At high doses, IKK16 is cytotoxic in a RIPK1 independent manner, and likely reflects off target effects of the inhibitor. Therefore, we cultured T cells in a range of IKK16 concentrations and assessed cell viability in the presence and absence of TNF. In the absence of CASPASE8 expression, both naive CD4 and naive CD8 T cells were resistant to FASL induced cell death observed in WT T cells from control mice (Fig. 5A). Similarly, while T cells from control mice were sensitised to TNF induced cell death in the presence of the IKK inhibitor IKK16, *Casp8* deficient T cells were completely resistant to TNF dependent cell death (Fig. 5B). Of incidental note, cytotoxicity of IKK16 at high doses, in the absence of TNF, which we have previously shown is RIPK1 independent (Carty et al., 2023), also appeared to be CASPASE8 independent, since similar profile of cell death was observed in the absence of CASPASE8 expression. Thus, the cytotoxic effects of IKK16 do appear to be off target effects independent of extrinsic cell death pathways. In conclusion, the naive T cells present in Casp8ΔT^CD2^ mice appear resistant to both cell death inducing stimuli with no evidence of either apoptotic or necroptotic cell death.

**Figure 5.**
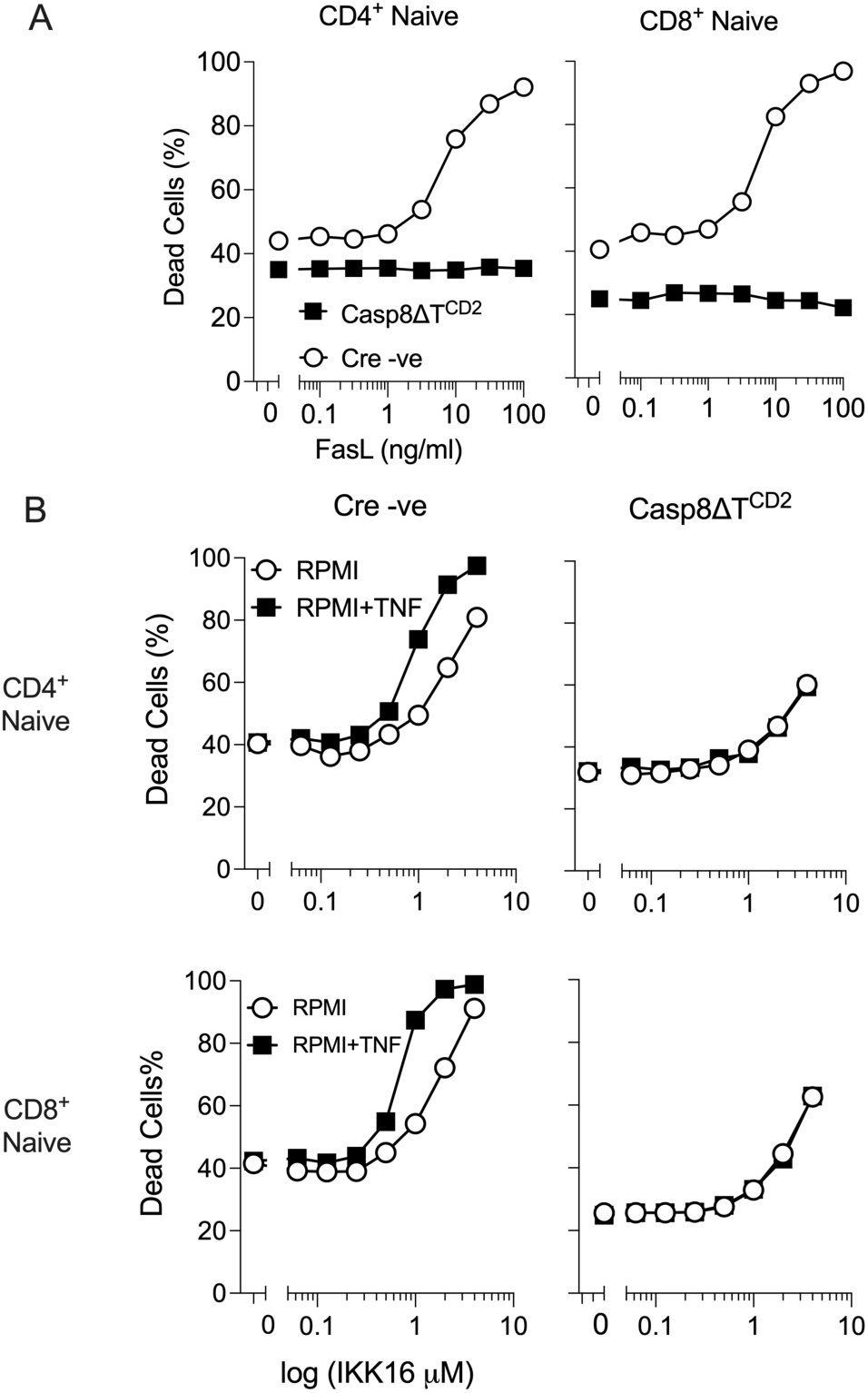
CASPASE8 deficient T cells are resistant to TNFRSF induced death. (A) T cells from Casp8ΔT^CD2^ mice or Cre-ve littermates were cultured overnight with a titration of FASL and crosslinking reagent, and then cell viability of naive CD4^+^ and CD8^+^ T cells assessed using a Live/Dead dye by flow cytometry. (B) T cells were cultured with a titration of IKK16 inhibitor in the presence and absence of TNF. Graphs show cell death of the indicated naive T cells subsets (rows) from the indicated mouse genotype (columns) with increasing IKK16 concentrations in the presence or absence of TNF. Data are representative of three independent experiments.

### CASPASE8 dependent survival of T cells is only in part dependent upon RIPK1 kinase activity

Culturing T cells from Casp8ΔT^CD2^ mice with TNF and FAS ligand in vitro did not reveal any evidence that CASPASE8 deficient T cells were sensitised to any form of cell death in the absence of CASPASE8 that might account for loss of T cells in vivo, while T cells from Casp8ΔT^CD2^ mice exhibited apparently normal lifespan. However, induced deletion of *Casp8* in peripheral CD8 T cells resulted in a rapid loss of a fraction of the peripheral CD8 compartment, so it was possible that those T cells that remain in adult Casp8ΔT^CD2^ mice that are resistant to cell death may not be fully representative of a replete CD8 T cell compartment. Since CASPASE8 has a well characterised role in repressing necroptosis, we wished to directly test, genetically, whether necroptotic cell death could account for any phenotypes in Casp8ΔT^CD2^ mice. We did this by breeding and analysing Casp8ΔT^CD2^ mice expressing a kinase dead RIPK1^D138N^ mutant (Newton et al., 2014). RIPK1 is a critical adaptor, triggered by its autophosphorylation, to promote formation of the necroptosome and induce necroptosis following TNFRSF signalling (Tenev et al., 2011). Therefore, kinase dead RIPK1 is anticipated to block necroptotic cell death in Casp8ΔT^CD2^ mice. Our prior analysis shows that lose of RIPK1 kinase activity does not impact thymic development (Blanchett et al., 2022). We first analysed the thymus of Casp8ΔT^CD2^ RIPK1^D138N^ mice to see whether the reduced thymus size and absence of mature NKT cells was rescued by RIPK1^D138N^. Comparing Cre –ve and Cre +ve littermates from Casp8ΔT^CD2^ RIPK1^D138N^ mice showed that numbers of both DN4 and DP thymocytes were similar in both cases (Fig. 6A), as were all other major thymic subsets (Fig. S2A). In contrast, mature NKT cell numbers in the thymus were subject to a very limited but detectable rescue when RIPK1 kinase was inactive (Fig. 6B and fig. S2B). Therefore, we found evidence of necroptosis amongst thymocytes following CASPASE8 ablation that modulated thymic development. CD4^+^ T cell numbers were normal in Casp8ΔT^CD2^ mice, so we next focused attention on the peripheral CD8 compartment in Casp8ΔT^CD2^ RIPK1^D138N^ mice. In Casp8ΔT^CD2^ mice, there were significant reductions in naive, CM and VM CD8^+^ subsets and a non-significant increase in EM (Fig. 2). In Casp8ΔT^CD2^ RIPK1^D138N^ mice, numbers of naive and CM subsets were similar between Cre –ve and Cre +ve littermates, while VM compartment was only reduced to around half that of Cre –ve control mice, suggesting RIPK1^D138N^ mediated partial rescue of this population (Fig. 6C). The CD8 EM compartment was more robustly expanded in Casp8ΔT^CD2^ RIPK1^D138N^ mice (Fig. 6C) than observed in Casp8ΔT^CD2^ mice (Fig. 2).

**Figure 6.**
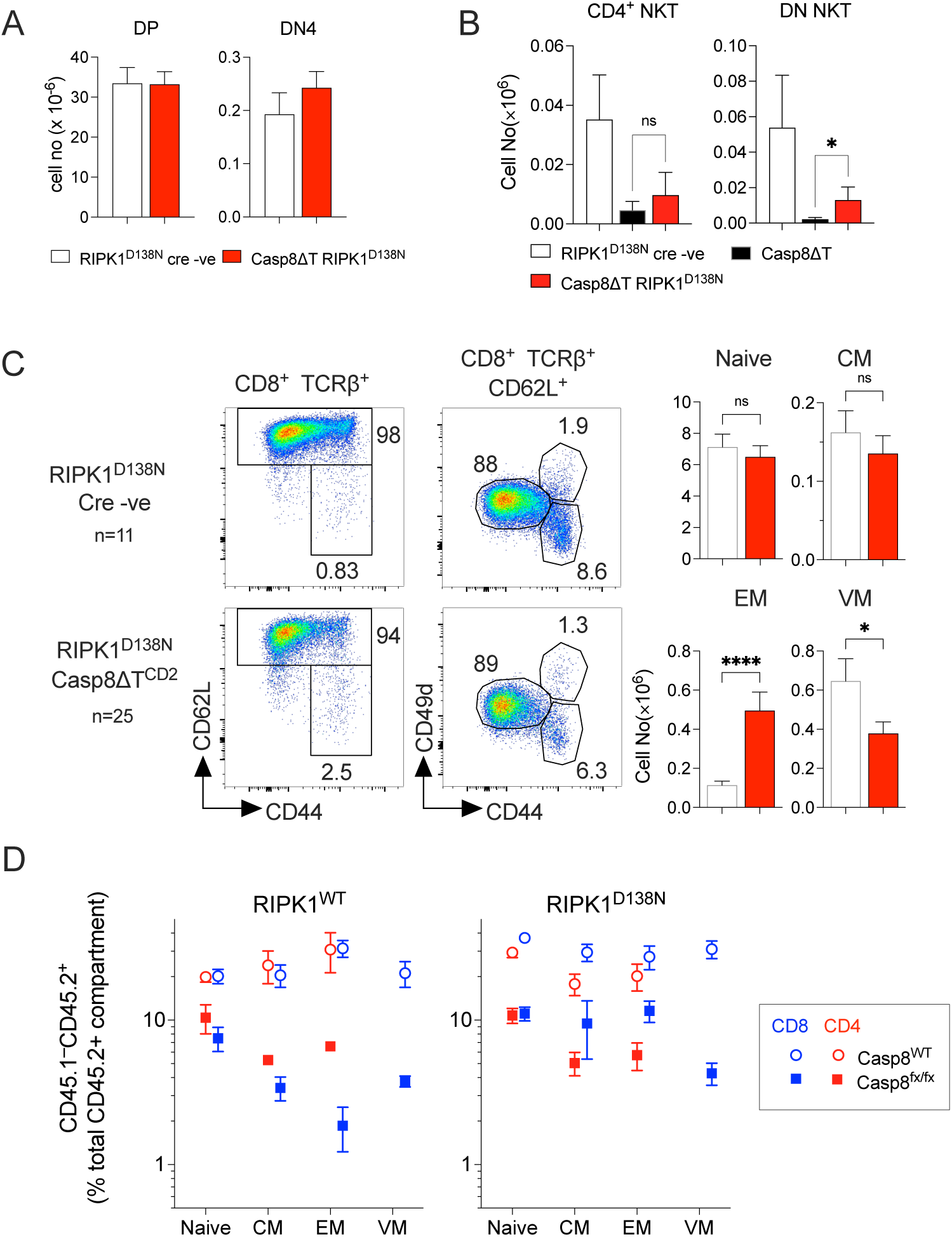
Partial rescue of T cell deficiencies in *Casp8* deficient mice by kinase dead RIPK1. Lymphoid organs (thymus, LN and spleen) from Casp8ΔT^CD2^ RIPK1^D138N^ (n=25) and Cre –ve littermates (n=11) were enumerated analysed by flow. (A) Bar charts show numbers of CD4^+^CD8^+^ DP and DN4 subsets in thymus of the indicated mice. (B) Bar charts are of numbers of CD1d-α-GalCer binding CD4^+^ and DN NKT cells in thymus of the indicated strains. (C) Density plots are of CD62L vs CD44 by TCRβ^+^CD8^+^ lymph node cells and show gates used to identify CD8 EM and CD62L^hi^ CD8^+^TCRβ^+^ T cells. Density plots of CD49d vs CD44 by CD62L^hi^ CD8^+^TCRβ^+^ T cell show gates used to identify naive, CM and VM subsets. Bar charts are of total numbers of the indicated subsets of CD8^+^ T cells recovered from both spleen and lymph nodes of the indicated strains. (D) Mixed bone marrow chimeras were generated by reconstituting CD45.1 hosts (n=4 per condition) with mixture of bone marrow from CD45.2 CD45.1 WT congenic donors and bone marrow from Cre + or Cre –ve donors of either Casp8ΔT^CD2^ or Casp8ΔT^CD2^ RIPK1^D138N^ donors (both CD45.2) (see materials and methods). 8 weeks later the composition of CD45.2 donor compartments was measured. Dot plots show the contribution of Casp8WT and Casp8 deficient cells to the indicated subsets of CD4 and CD8 T cells, originating from either RIPK1WT donors (top) or RIPK1 ^D138N^ expressing donors, as a fraction of the total donor CD45.2 compartment. Data are representative of three independent experiments.

Analysis of intact Casp8ΔT^CD2^ RIPK1^D138N^ mice provided evidence that necroptosis could account for some but not all of the phenotypes observed in Casp8ΔT^CD2^ mice. However, because the mutant RIPK1^D138N^ is expressed in all cells of these hosts, we wished to confirm the cell intrinsic nature of the rescue by generating mixed bone marrow chimeras using normal hosts. Irradiated CD45.1 hosts were reconstituted with a mixture of control bone marrow from CD45.1 CD45.2 WT donors, and bone marrow from either Cre + or Cre –ve donors from Casp8ΔT^CD2^ strain mice or Casp8ΔT^CD2^ RIPK1^D138N^ mice. Hosts were allowed to reconstitute for 8 weeks, and then the donor composition of CD4 and CD8 T cell compartments analysed across naive, CM, EM and VM subsets. As expected, donor reconstitution, relative to CD45.1 CD45.2 WT competitor, by bone marrow from WT Cre –ve littermates of Casp8ΔT^CD2^ mice exhibited a consistent and stable contribution across different subsets of both CD4 and CD8 T cells (Fig. 6D). Donor bone marrow from Casp8ΔT^CD2^ mice reconstituted CD8 compartments poorly, with a low level of competitive chimerism amongst naive CD8 T cells that was progressively reduced in EM, CM and VM subsets. Analysing chimerism in CD4 subsets revealed a similar behaviour, with a reduced representation of *Casp8* deficient T cells across all CD4 subsets analysed. We compared this with chimerism from chimeras using donor bone marrow from RIPK1^D138N^ mice. Cre –ve bone marrow from RIPK1^D138N^ strains also exhibited a consistent and stable contribution across of different subsets of both CD4 and CD8 T cells. Amongst CD8 subsets, kinase dead RIPK1 did not improve the competitive fitness of *Casp8* deficient naive or VM T cells, which were as equally under represented as the same subsets in chimeras using bone marrow from Casp8ΔT^CD2^ mice. However, donor representation amongst CM and EM subsets remained stable, suggesting RIPK1^D138N^ did prevent further out-competition (Fig. 6D). Amongst CD4 subsets, kinase dead RIPK1 did not rescue out-competition of naive, or CM and EM subsets which were further reduced in representation. These data reveal that competitive fitness of T cells is reduced in the absence of CASPASE8 expression, for both CD8 and CD4 subsets, and that blocking RIPK1 kinase activity only rescues some subsets, notably CD8 EM and CD8 CM.

## Discussion

In the present study, we wished to better understand the impact and mechanisms by which CASPASE8 signaling influences the development and maintenance of the T cell compartments of mice. Analysing the chief developmental and mature compartments revealed important roles for CASPASE8 at multiple stages of differentiation and in different lineages. Analysing mice with kinase dead RIPK1 revealed that only some of the observed T cell defects in the absence of CASPASE8 could be accounted for by a loss of cell death control and induction of necroptosis. In order to reconcile the range and complexity of phenotypes we observed, we are inevitably led to the conclusion that CASPASE8 must promote T cell survival by additional mechanisms to just those dependent on repressing necroptosis and that the phenotype of Casp8ΔT^CD2^ mice is therefore the result of a complex, compound phenotype.

While it is clear that activated T cells require CASPASE8 to protect them from induction of necroptosis (Ch’en et al., 2008;Ch’en et al., 2011;Feng et al., 2019), previous studies have been somewhat equivocal regarding the requirement for CASPASE8 to establish and/or maintain normal homeostasis of mature peripheral T cell compartments (Salmena et al., 2003;Ch’en et al., 2008;Ch’en et al., 2011). Our detailed analysis revealed defects amongst DN thymocytes, CD8 CM, VM and NKT cells that had not previously been recognised. In regard to naive T cells, however, evidence appeared somewhat conflicting, with reports of both normal compartment size (Ch’en et al., 2008) and either selective (Ch’en et al., 2011) or broad T cell lymphopenia (Salmena et al., 2003). In Casp8ΔT^CD2^ mice, we observed a clear reduction in the size of the naive CD8 T cell compartment while, numbers of naive CD4 T cells appeared normal. We did find evidence for a modest impact on thymic development, due to RIPK1 dependent necroptosis amongst DN progenitors. In contrast, post selection SP thymocyte numbers were largely normal, and so it appears that the modest reduction in DP cellularity may be compensated by more efficient positive selection to maintain normal SP compartments and thymic output, and therefore it does not appear that a deficiency in post selection thymic precursors can account for the substantial reduction in the CD8 T cell compartment.

Amongst peripheral T cells, life spans and turnover of T cells from Casp8ΔT^CD2^ mice appeared normal, so it did not appear that compartment wide alteration of homeostatic properties of the CD8 T cells could account for the observed reduction in compartment size. Evidence that a subset of naive T cells may be specifically and acutely dependent on continued CASPASE8 expression for their survival, came from experiments revealing the acute loss of a fraction of CD8 T cells following induced *Casp8* deletion amongst mature CD8 T cells. In the steady state, such CASPASE8 dependent CD8 T cells in Casp8ΔT^CD2^ mice may be rapidly purged as they are generated, such that the peripheral compartment is reduced in size, but that remaining cells have otherwise normal life spans, even when transferred to a WT host environment. Kinase dead RIPK1 appeared to restore the naive CD8 compartment in Casp8ΔT^CD2^ mice, suggesting that the loss of some naive CD8 T cells was due to necroptosis. This is surprising because neither thymocytes nor naive T cells express *Mlkl*, that is critical for induction of necroptosis (Webb et al., 2019). These observations could be reconciled if *Mlkl* expression was induced transiently in some CD8 T cells, possibly during development or maturation, that could render them briefly susceptible to cell death in the absence of CASPASE8, such that detection of *Mlkl* expression amongst bulk CD8 T cells remain hard to detect. Such a possibility is supported by the observations that type I interferons are reported to induce *Mlkl* in other cell types (Knuth et al., 2019), and developing thymocytes do express an IFN-induced gene signature during development (Xing et al., 2016). Whether such a mechanism is active in CD8 T cells remains to be determined however.

Our studies found evidence of more than one survival mechanism controlled by CASPASE8. We analysed mice with kinase dead RIPK1 to identify those phenotypes that were RIPK1 kinase dependent and may result from induction of necroptosis. Thymic cellularity and peripheral naive CD8 T cell numbers were rescued by RIPK1^D138N^ as was generation of CD8 CM and CD8 EM in both intact mice and in bone marrow chimeras. These findings are consistent with previous studies demonstrating that effector T cells are susceptible to necroptosis in the absence of CASPASE8, both following activation in vitro and during viral infection in vivo (Ch’en et al., 2008;Ch’en et al., 2011;Feng et al., 2019). In contrast, CASPASE8 deficient CD8 VM T cells were only partially rescued by kinase dead RIPK1, while the profound loss of NKT cells was barely impacted at all by inactivation of RIPK1. It was also striking that CASPASE8 deficient T cells were highly resistant to cell death induction in vitro, in response to FAS and TNF stimuli, demonstrating that resting T cells are not obviously sensitised to necroptosis in the absence of CASPASE8, at least in response to TNFRSF stimuli. Further evidence of a CASPASE8 dependent survival mechanism that did not depend on repression of necroptosis came in the mixed bone marrow chimeras, where there were clear defects in the generation and/or persistence of CD4 CM and CD4 EM, that were not rescued by kinase dead RIPK1. Also, naive CD4 and naive CD8 T cells were both underrepresented in mixed bone marrow chimeras reconstituted with donors expressing RIPK1^D138N^, even though kinase dead RIPK1 rescued normal thymic development.

While CASPASE8 is a potent trigger of cell death, our findings suggest that another important role of CASPASE8 in T cells is to promote their survival. This is in part because of the important role CASPASE8 plays in repressing necroptosis, as was evident in developing thymocytes and in CD8 CM and CD8 EM. Indeed, when necroptosis was blocked in RIPK1^D138N^ mice, CASPASE8 deficiency resulted in a substantial expansion of CD8 EM compartment, that resembles similar expansions observed in *Fas^lpr^* mice (Giese and Davidson, 1992;Russell et al., 1993), and likely represents a failure of FAS to kill CD8 EM in the absence of CASPASE8 and RIPK1 kinase activity, as also described in FADD and RIPK1 deficient mice (Zhang et al., 2011). However, there was also abundant evidence for a pro-survival function of CASPASE8 that was distinct from its role in repressing necroptosis. This function appears to be important in NKT cells and CD8 VM, and may also be required for optimal competitive fitness of naive CD4 and CD8 T cells. The mechanism for this survival function remains obscure. We could not find any evidence that well characterised survival signals that derive from TCR recognition of self-peptide MHC complexes or from IL-7R signalling were affected in the absence of CASPASE8. Oligomerisation of CASPASE8 is associated with triggering apoptosis and active CASPASE8 is also responsible for targeted cleavage of RIPK1 (Newton et al., 2019;Newton et al., 2024). It is possible that CASPASE8 has other substrates whose cleavage is required for normal T cell survival. Alternatively, functions have been described for CASPASE8 that are independent of its proteolytic activity. Studies of TRAIL signalling in cell lines show that CASPASE8 has an adapter function during signalling that is required for NF-κB dependent inflammatory signalling and induction of cytokine production downstream of TRAIL receptor, that does not require enzymatic activity by CASPASE8 (Henry and Martin, 2017). Acute T cell survival is dependent upon tonic NF-κB signalling (Carty et al., 2023), and so it is possible CASPASE8 may be required for transmission of these signals. To explore these possibilities, future studies will need to address whether enzymatic or adapter function of CASPASE8 are required for RIPK1 independent survival mechanisms.

## Materials and methods

### Mice

Mice with the following mutations were used in this study; conditional alleles of Casp8, B6.129-Casp8^tm1Hed^/J (*Casp8^flox^*)*, Cre* transgenes expressed under the control of the human CD2, B6.Cg-Tg(CD2-icre)4Kio/J (*huCD2^iCre^*) (de Boer et al., 2003) mice with a D138N mutation in *Ripk1, B6.129-Ripk1^tm1Geno^/J* (RIPK1^D138N^) (Newton et al., 2014) and Tg(Cd8a-icre/ERT2,-GFP)Daav (CD8^CreERT2^) (Andrews et al., 2021). The following strains were bred for this study; *Casp8^fx/fx^ huCD2^iCre^* (Casp8ΔT^CD2^), *Casp8^fx/fx^ huCD2^iCre^* RIPK1^D138N^ (Casp8ΔT^CD2^RIPK1^D138N^), *Casp8^fx/fx^ CD8^CreERT^,* SJL.C57Bl6/J (**CD45.1**), (SJL.C57Bl6/J x C57Bl6/J)F1 (CD45.1 CD45.2). Cre activity in CD8CreERT mice was induced *in vivo* by i.p. injection of 2mg of tamoxifen in corn oil for five consecutive days. All mice were bred in the Comparative Biology Unit of the Royal Free UCL campus and at Charles River laboratories, Manston, UK. Animal experiments were performed according to institutional guidelines and Home Office regulations.

Mixed bone marrow chimeras were generated by irradiating CD45.1 C57Bl6/J hosts with 800 RADS, followed by injection with total 10^7^ T depleted bone marrow cells. Bone marrow was isolated from WT CD45.1 CD45.2 C57Bl6/J donors, and from Casp8ΔT^CD2^ and Casp8ΔT^CD2^ RIPK1^D138N^ strain mice that are both CD45.2. Bone marrow from CD45.1 CD45.2 donors was mixed 1:1 with either Cre –ve or Cre +ve donors from the different *Casp8* strains.

### Flow cytometry and electronic gating strategies

Flow cytometric analysis was performed with 4-5 × 10^6^ thymocytes, lymph node or spleen cells. Cell concentrations of thymocytes, lymph node and spleen cells were determined with a Scharf Instruments Casy Counter. Cells were incubated with saturating concentrations of antibodies in 100 μl of Dulbecco’s phosphate-buffered saline (PBS) containing bovine serum albumin (BSA, 0.1%) for 1hour at 4°C followed by two washes in PBS-BSA. Panels used the following mAb: PE-Cy7-conjugated antibody against CD25 (eBioscience), PE-conjugated antibody against CD127 (ThermoFisher Scientific), BV785-conjugated CD44 antibody (Biolegend), BV421-conjugated antibody against CD4 (Biolegend), BUV395-conjugated antibody against CD8 (BD Biosciences), BV711-conjugated antibody against CD24 (BD Biosciences), PerCP-Cy5.5-conjugated antibody against TCR (Cambridge Biosciences), FITC-conjugated antibody against CD5 (eBiosciences), BV650-conjugated antibody against CD45.1 (Biolegend), FITC-conjugated antibody against CD45.2 (ThermoFisher Scientific), APC-conjugated antibody against CD49d (Biolegend), FITC-conjugated antibody against Ki67 (ThermoFisher Scientific). Cell viability was determined using LIVE/DEAD cell stain kit (Invitrogen Molecular Probes), following the manufacturer’s protocol. multi-colour flow cytometric staining was analysed on a LSRFortessa (Becton Dickinson) instrument, and data analysis and colour compensations were performed with FlowJo V10 software (TreeStar). The following gating strategies were used: peripheral CD4 naive T cells - CD4^+^ TCRβ^+^CD25^−^CD44^lo^CD62L^hi^, CD4 central memory (CM) T cells - CD4^+^ TCRβ^+^CD25^−^ CD44^hi^CD62L^hi^, CD4 effector memory (EM) T cells - CD4^+^ TCRβ^+^CD25^−^CD44^hi^CD62L^lo^, CD8 naive T cells - CD8^+^ TCRβ^+^CD44^lo^CD62L^hi^CD49d^lo^, CD8 CM - CD8^+^ TCRβ^+^CD44^hi^CD62L^hi^CD49d^hi^, CD8 virtual memory (VM) T cells - CD8^+^ TCRβ^+^CD44^hi^CD62L^hi^CD49d^lo^ and CD8 EM - CD8^+^ TCRβ^+^CD44^hi^CD62L^lo^. Immature and mature CD4^+^ SP thymocytes were identified as CD4^+^ CD8^−^ TCRβ^+^HSA^hi^CD62L^lo^ and CD4^+^ CD8^−^TCRβ^+^HSA^lo^CD62L^hi^ respectively. Immature and mature CD8^+^ SP thymocytes were identified as CD4^−^ CD8^+^TCRβ^+^HSA^hi^CD62L^lo^ and CD4^−^ CD8^+^TCRβ^+^HSA^lo^CD62L^hi^ respectively.

### In vitro culture

Lymph node T cells were cultured at 37°C with 5% CO2 in RPMI-1640 (Gibco, Invitrogen Corporation, CA) supplemented with 10% (v/v) fetal bovine serum (FBS) (Gibco Invitrogen), 0.1% (v/v) 2-mercaptoethanol βME (Sigma Aldrich) and 1% (v/v) penicillin-streptomycin (Gibco Invitrogen) (RPMI-10). Recombinant TNF was supplemented to cultures at 20ng/ml, unless otherwise stated, and was obtained from Peprotech, with PBS used as vehicle. Recombinant FASL (Enzo Life Sciences UK Ltd.) was supplemented to cultures at indicated concentrations together with fixed concentration (10 μg/ml) of cross-linking enhancer. Inhibitors were used at the following concentrations, unless otherwise stated: IKK16 (2µM in 0.1% DMSO).

### Statistics

Statistical analysis, line fitting, regression analysis, and figure preparation were performed using Graphpad Prism 8. Column data compared by unpaired Mann-Witney student’s t test. ns = nonsignificant, * p<0.05, ** p<0.01, *** p<0.001, **** p<0.0001.

## Non-standard abbreviation list

CBM: CARD11 BCL10 MALT1
CIAP: Cellular inhibitor of apoptosis protein
IKK: Inhibitor of kappa B kinase
IκB: Inhibitors of kappa B
Nec1: Necrostatin-1
RIPK1: Receptor-interacting protein kinase 1
TAK1: Transforming growth factor-beta activated kinase 1
TNFR1: Tumour necrosis factor receptor 1
TRADD: Tumour necrosis factor receptor 1-associated death domain protein
TRAF: Tumour necrosis factor receptor-associated factor

## Acknowledgements

We thank UCL Comparative Biology Unit staff for assistance with mouse breeding and maintenance. We thank the following for generously sharing of their mouse strains: Prof Vishva Dixit for the RIPK1^D138N^ strain. The authors declare no competing financial interests. The work in the Seddon lab is supported by the Medical Research Council UK under programme codes MR/ P011225/1.

