## Supplementary figures for "Evidence for a RIPK1-independent survival mechanism for CASPASE-8 in αβ T cells"

**Figure S1 - Limited Cre toxicity amongst CD8 EM and CD8 VM in CD8$^{CreERT}$ mice.** *CD8$^{CreERT}$* mice were treated with TAM for five consecutive days. At days 7 and 21, mice were culled and CD8 T cell compartment in lymph node and spleen enumerated. (A) Density plots show representative phenotypes in lymph nodes cells from TAM treated CD8$^{CreERT}$ mice (n=5), untreated CD8$^{CreERT}$ mice (n=4) and Cre -ve littermate controls (n=5) at day 21, and gates used to identify naive, EM, CM and VM subsets of CD8 T cells. (B) Bar charts show total numbers of the indicated CD8 subset recovered from lymph node and spleen of the indicated strains. Data are representative of 2 independent experiments.

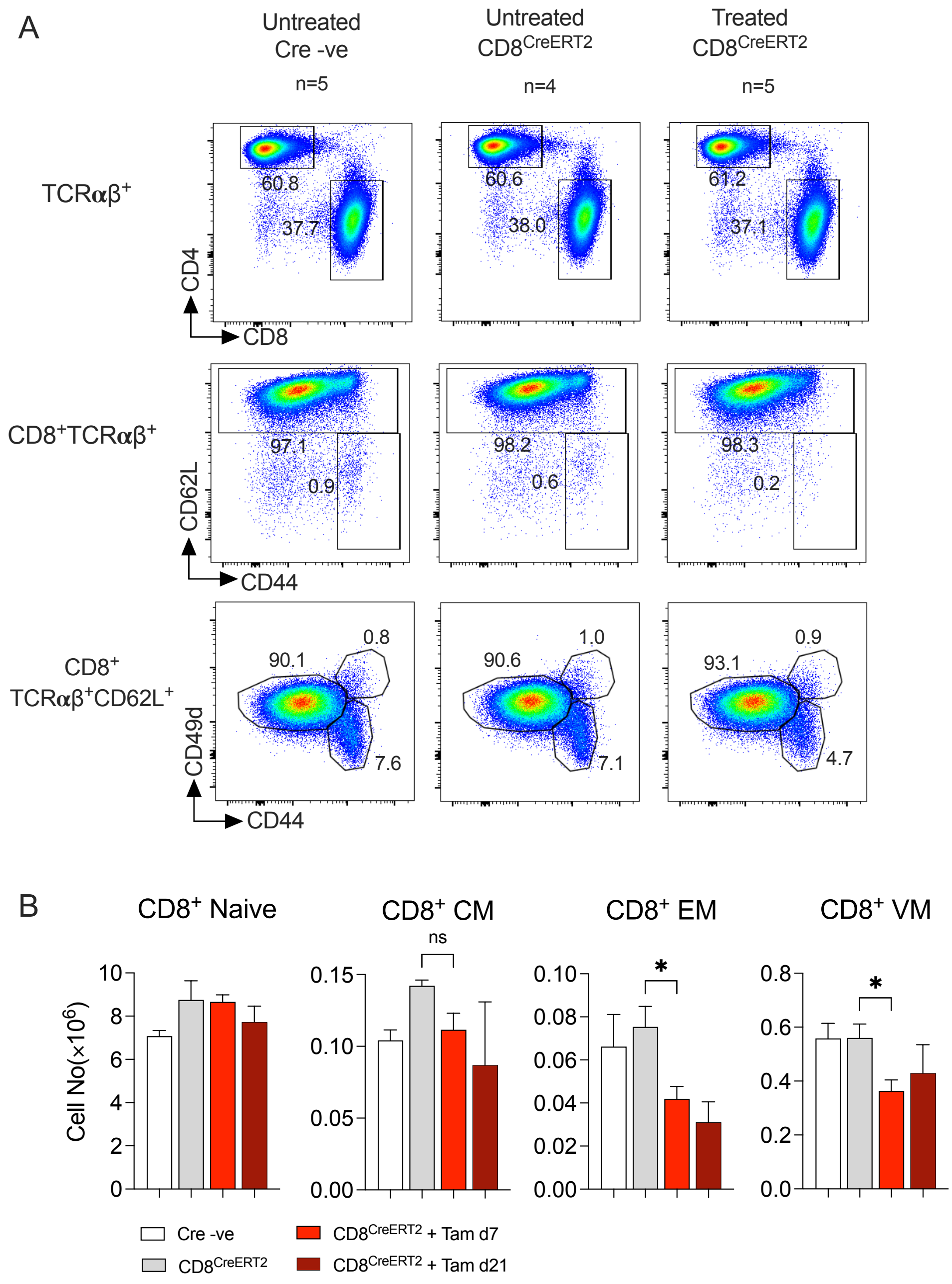

**Figure S2 - Partial rescue of thymopoiesis in Casp8ΔT$^{CD2}$ mice by kinase dead RIPK1.** Lymphoid organs (thymus, LN and spleen) from Casp8ΔT$^{CD2}$ RIPK1$^{D138N}$ (n=25) and Cre –ve littermates (n=11) were enumerated analysed by flow. (A) Bar charts show numbers of the indicated thymic subsets from Casp8ΔT$^{CD2}$ RIPK1$^{D138N}$ mice and Cre –ve littermates. (B) Density plots are of HSA vs CD1d-α-GalCer (CD1dTet) binding to total live thymocytes. CD4 vs CD8 expression is for CD1dTet$^{+}$ cells. HSA vs CD44 expression is shown for CD4$^{+}$ CD8$^{-}$ CD1dTet$^{+}$ (CD4 NKT) cells and CD4$^{-}$CD8$^{-}$ CD1dTet$^{+}$ (DN NKT) cells. Bar charts summarise total numbers of CD4$^{+}$ NKT and DN NKT recovered from the indicated strains.

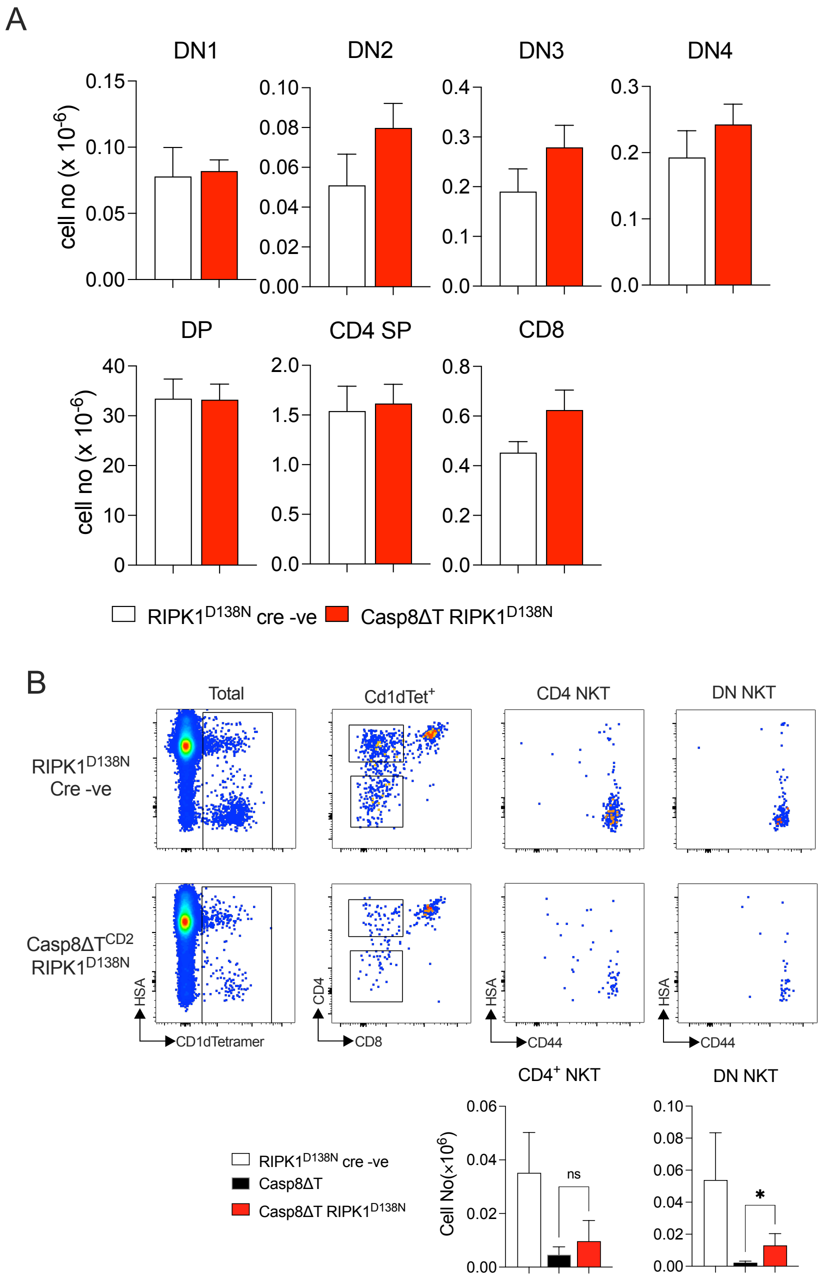
